# Neural Synchrony in Parent-Child Dyads: Profiles Associated with Interparental Conflict and Internalizing Symptoms

**DOI:** 10.1101/2025.07.02.662817

**Authors:** Charles Alvarado, Carlomagno C. Panlilio, Koraly Perez-Edgar, Khalil I. Thompson, Denny Schaedig, Nadine Melhem, Susan B. Perlman

## Abstract

Interparental conflict and parental stress are well-established risk factors for child psychopathology, including elevated internalizing and externalizing symptoms. From a family systems framework, these stressors may spill over into the parent-child relationship, undermining emotional attachment and co-regulation processes central to children’s mental health. Neural synchrony, defined as the dynamic, mutual alignment of brain activity between a parent and child, offers a biological index of these dyadic processes. Using functional near-infrared spectroscopy, researchers have shown that greater neural synchrony (NS) in prefrontal brain regions is associated with more attuned caregiving and positive child adjustment. Yet, NS is not uniform; it varies across dyads in pattern and regional distribution, potentially reflecting differences in relational dynamics, regulation, or stress exposure. To capture this heterogeneity, we used latent profile analysis to identify distinct synchrony patterns along the right and left ventrolateral and dorsolateral prefrontal cortices during the DB-DOS:Biosynch – a mild stress, three-context task. We further examined whether interparental conflict and perceived parental stress predicted profile membership, and whether children’s internalizing and externalizing behaviors differed by profile. Among 194 dyads, two profiles emerged: lower baseline synchrony (LB; *n* = 132) and higher baseline synchrony (HB; *n* = 62). Greater interparental conflict reduced the odds of membership in HB, while parental stress was not predictive of profile membership. Additionally, children in LB exhibited higher levels of internalizing behaviors compared to HB, with no group differences observed for externalizing behaviors. These findings underscore the value of capturing synchrony heterogeneity in understanding family stress and child psychopathology.

## Introduction

Conflict is a normative part of relationships within family systems where children observe disagreements between parents. However, when interparental conflict is frequent, intense, or poorly resolved, it can have deleterious effects (Van Eldik et al., 2020). Consequently, children exposed to such conflicts are at heightened risk for difficulties in emotion regulation, physiological stress reactivity, and social functioning (Davies et al., 2006; Harold & Sellers, 2018). Although these associations are well-documented, the mechanisms through which interparental conflict influences children’s development remain less understood.

To better understand these connections, it is useful to view the family as a network of interconnected subsystems as conceptualized by family systems theory (FST) (Cox & Paley, 1997; Harold & Sellers, 2018). FST’s spillover hypothesis states that disruptions can transfer from the interparental subsystem to the parent-child subsystem, impairing emotional attunement, co-regulation, and parenting practices (Fosco & Grych, 2007; Gao et al., 2019). Children absorb these disruptions both directly and indirectly, with even low-quality interparental relationships, linked to elevated internalizing and externalizing symptoms over time (Davies et al., 2006; Van Eldik et al., 2020). One candidate mechanism that links these relational stressors to children’s outcomes is parent-child synchrony, which may offer insight into how such stress “gets under the skin” (Gray et al., 2024).

*Parent-child synchrony*, the moment-to-moment coordination of affective, behavioral, and physiological states between a caregiver and child (Birk et al., 2022), is associated with child adjustment and stress resilience (Davis et al., 2018; Feldman, 2007, 2015; Quiñones-Camacho et al., 2022). Behavioral and biological synchrony, such as shared gaze (e.g., Beebe et al., 2016), mutual affect (e.g., Vanoncini et al., 2022), and aligned autonomic arousal (e.g., Lunkenheimer et al., 2021), support secure attachment and healthy development (Feldman, 2007; Meins et al., 2012). Synchrony interactions are dynamic, shaped by each partner’s regulatory capacities, stress exposure, and relational history (Feldman, 2015; Kim et al., 2010). These patterns, whether consistently high, low, or variably in sync, influence how dyads navigate emotionally charged moments. Examining synchrony in daily contexts allows researchers to capture how residual stress alters the quality of dyadic engagement (Birk et al., 2022; van Dijk et al., 2022).

A more recent approach, neural synchrony (NS) – the coordinated brain activity between parent and child – has emerged as a promising index of shared attention, affective alignment, and mutual engagement (Nguyen et al., 2024; Quiñones-Camacho et al., 2020). Using hyperscanning (i.e., the measurement of two individuals’ brain activity simultaneously), researchers have demonstrated that greater parent-child NS, particularly in the frontal cortex, is associated with more attuned caregiving and better child outcomes (Bi et al., 2023; Reindl et al., 2018).

Functional near-infrared spectroscopy (fNRIS) has become increasingly applied in this research area due to its non-invasive, motion-tolerant, and developmentally appropriate design (Cui et al., 2012; Quiñones-Camacho et al., 2020). Studies using fNIRS have linked prefrontal synchrony to children’s emotion regulation, cooperative problem-solving, and engagement during joint tasks (Nguyen et al., 2024; Reindl et al., 2018). In addition, environmental contexts such as family stress reduce parent-child NS (Azhari et al., 2019; Hoyniak et al., 2021). These findings raise important questions about the mechanisms by which relational stress becomes biologically embedded, suggesting the need to identify variation in how synchrony manifests across different family contexts.

The current study uses a novel application of latent profile analysis (LPA) to examine heterogeneity in parent-child synchrony across global and lateralized (i.e., spatially specific) prefrontal regions, with the goal of identifying subgroups that differ in relation to interparental conflict, family stress, and child adjustment outcomes. This approach extends prior work by capturing meaningful variation in parent-child NS. The aim is to explore whether distinct patterns of synchrony reflect differential processes within the dyad, potentially shaped by the family stress context, and offer insight into heterogeneity in children’s adjustment outcomes.

## Method

### Participants

The data were drawn from the Child Affect and Resilience to Experiences (CARE) study, a longitudinal investigation examining how parental conflict influences the development of biological stress and its links to early childhood psychopathology over time. The CARE sample included 229 children ages 4 to 7 years (114 male; *M* = 5.74 years, *Range* = 4–8 years).

According to parent report, 69% of children were White, 12% Black, 1% Asian, and 18% multiracial; 11% identified as Hispanic/Latinx. Among 194 participating parents, 77% self-identified as White, 12% as Black, 6% as Asian, <1% as Pacific Islander, and 4% as multiracial; 8% identified as Hispanic/Latinx. Most parents were female (92%), and multiple children per family were included in the sample.

In order to enrich the sample for parental conflict, letters were sent to divorcing families through publicly available divorce records in local counties. We also recruited through Kids in the Middle, a local clinic treating divorcing families. Finally, families were recruited through local advertising. Exclusion criteria included head trauma with loss of consciousness; developmental disability that could impair task comprehension (e.g., autism spectrum disorder); exposure to major stressors (e.g., parental death, incarceration, or serious illness); chronic illness affecting stress systems or had recently used corticosteroids or other medications influencing neuroendocrine function in the child. Procedures for the current study were approved by the Washington University in St. Louis Institutional Review Board.

## Questionnaires

### Interparental Conflict

For analytic purposes, interparental conflict was dichotomized to reflect low versus high conflict. Interparental conflict was classified as high if parents met at least one of the following criteria: 1) a total Revised Dyadic Adjustment Scale (RDAS) score below the recommended cutoff of 48 (Anderson et al., 2014), or 2) parental separation within the past year. The RDAS is a 14-item instrument that assesses romantic relationship quality across three domains: consensus, satisfaction, and cohesion(Anderson et al., 2014). Items are rated on a six-point Likert-type scale, with lower scores indicating poorer relationship quality and greater interparental conflict (*α* = 0.998).

### Perceived Parental Stress Scale

The Perceived Parental Stress Scale (PSS; Berry & Jones, 1995) is a caregiver-report measure designed to assess parents’ subjective experiences of stress and satisfaction in response to parenting demands. It captures dimensions such as emotional strain, role overload, and the challenges of daily caregiving. The 10-items used in this study are rated on a five-point Likert-type scale, ranging from 1 (strongly disagree) to 5 (strongly agree), with higher scores reflecting greater perceived parenting stress. A continuous total score was used in these analyses (*α* = 0.400).

### Internalizing and Externalizing Symptoms

The MacArthur Health Behavior Questionnaire (HBQ; Armstrong et al., 2003) is a comprehensive caregiver-report tool designed to assess comorbid psychiatric symptoms, functional impairment, and treatment needs in children ages 4–8, offering a developmentally sensitive and integrative approach to understanding early psychopathology. Though the HBQ assesses a broad range of child psychopathology symptoms, the present analyses focused specifically on the internalizing and externalizing domains. The internalizing domain combines items related to depression, overanxious, and separation anxiety subscales, while the externalizing domain combines items related to oppositional defiance, conduct problems, overt hostility, and relational aggression subscales. HBQ items for the internalizing and externalizing scales are Likert-type scales with three points (e.g., “never or not true,” “sometimes true,” “often or very true”). Mean scores across respective subscales for each domain are used in these analyses (Internalizing: *α* = 0.820; Externalizing: *α* = 0.903).

### fNIRS Dyadic Interaction Paradigm

The DB-DOS: BioSync paradigm (Quiñones-Camacho et al., 2022; Quiñones-Camacho et al., 2020) is a neuroimaging-adapted version of the Disruptive Behavior Diagnostic Observation Schedule (DB-DOS; Wakschlag et al., 2008) structured to examine parent–child interactions across a range of affective and behavioral demands in a format suitable for biological measurement. Designed with fNIRS hyperscanning in mind, the task allows for the capture of brain activation from both parent and child during shared, ecologically valid experiences. The protocol includes three primary conditions or contexts – baseline, stress, and recovery – each context containing four 2-minute trials separated by brief 15-second intertrial intervals to optimize neural signal quality. In the “baseline” condition, dyads are engaged in a relaxed joint art activity using paper, stickers, and markers, offering a low-arousal context for baseline comparison. Mild stress is introduced in the “stress” condition, where parents and children are left alone at a table with enticing toys placed directly next to the child. The dyad is told that they are not allowed to touch the toys until their tasks are completed. The dyads are then instructed to solve tangram puzzles – geometric shape challenges that were intentionally too difficult for the child’s developmental level. Dyads are told that they will only receive a prize depending on the completion of an unknown number of puzzles. The time to solve each puzzle is reduced from a promised 2 minutes to just 1 minute and 45 seconds, and a countdown timer is prominently displayed to reinforce time pressure. Finally, the recovery condition allows the dyad to interact freely with the previously restricted toys in a less structured, low demand setting designed to mirror the "baseline” condition while facilitating a return to emotional equilibrium. This design enables the capture of fluctuations in NS under shifting social and regulatory demands.

### fNIRS Data Acquisition

Optical imaging data were collected noninvasively using a continuous-wave fNIRS system (NIRScout; NIRx Medical Technologies LLC, Glen Head, NY, USA). Light at two wavelengths (760 nm and 850 nm) was emitted from eight LED sources and detected by four photodiode sensors, producing 10 source-detector pairs per wavelength. Signals were recorded at a sampling rate of 7.81 Hz. The optodes were arranged on a neoprene cap, with each source-detector pair spaced 3 cm apart. Cap placement followed the international 10–20 EEG system for both parent and child participants, with sources positioned at Fp1, Fp2, AF3, AF4, F5 (duplicated), FC5, and FC6. This configuration provided coverage across regions of the prefrontal cortex, including frontopolar, dorsolateral, and ventrolateral areas. Probe placement was guided by the fNIRS Optodes’ Location Decider toolbox (Zimeo Morais et al., 2018), which registered optode positions to Brodmann areas based on the brain atlas by Garey (1999). To enhance signal quality, hair was manually separated beneath the sensors as needed to ensure proper contact with the scalp.

### fNIRS Preprocessing

As we have done in previous studies (Fishburn et al., 2018; Thompson et al., 2025), we define parent-child NS as the average of cross-wavelet transform coherence (WTC) between the dyad during a particular context and approximate it using task-based frequencies of interest (between .0095-.2 Hz; Nguyen et al., 2020). We calculate WTC separately for each of the three task conditions: baseline, stress, and recovery.

We preprocessed the data using the MNE-Python package (Gramfort, 2013). First, whole-trial raw fNIRS data were converted to raw optical density utilizing the modified Beer-Lambert Law (MBLL) to account for variations in photon pathlength and tissue absorption essential for detecting hemoglobin. Second, each channel was evaluated using the scalp-coupling index (Hernandez & Pollonini, 2020; Pollonini et al., 2016) to assess the quality of optode connection to participants scalps as the fNIRS scan was recorded. Channels with an SCI < 0.5 (scale between 0 and 1) were marked as invalid and removed from further analysis. If either the parent or child in any dyad had ten or more (of their 20) channels marked as invalid due to low SCI, the dyad was dropped from the analysis (7 dyads). This is a conservative data quality cut-off compared to the majority of the published hyperscanning literature (Nguyen et al., 2020; Reindl et al., 2019). Third, the temporal-derivative distribution repair (TDDR) algorithm was administered on a channel-wise basis to detect fluctuations outside of the expected hemodynamic range and regionally smooth artifacts like head motion. The modified optical density data was converted to hemoglobin concentration values using a partial-pathlength factor (PPF) set at 0.1 via the Beer-Lambert Law. Subsequently, the hemoglobin signal was bandpass-filtered with low-and high-pass frequency thresholds of 0.01 and 0.2 Hz to target the natural frequency-based oscillatory dynamics of task-related hemoglobin changes in the brain (Reindl et al., 2018).

We standardized epoch-related timings across participants, yielding trial lengths of 120s, 105s, and 120s, for the baseline, stress, and recovery conditions, respectively. The stress condition involved a deliberate reduction in block duration to induce a mild increase in stress.

Four repeated trials constituted one of the three conditions of the task, and the preprocessed fNIRS data was divided into epochs covering the entire fNIRS time series. We did not analyze HbR (deoxygenated hemoglobin) channels in this project, as HbO (oxygenated hemoglobin) is more relevant to task-based imaging studies (Blockley et al., 2013). We performed an additional channel-pruning step, where if the peak-to-peak differences across an epoch for any given HbO channel (max value of epoch minus min value of epoch) was greater than 200e-6, it was removed to minimize the presence of spike signal artifacts. The channels of each epoch were then standardized such that the mean average of the first 5 seconds of the epoch was considered a baseline HbO value, and this amount was subtracted from the entire signal. Finally, a linear detrend to reduce overall signal variation was applied for each epoch when loaded for analysis in the MNE protocol.

NS values were estimated for each parent and child (i.e., child of Dyad A paired with parent of Dyad A; a “real dyad”) in a task condition by fNIRS channel analysis. Prior to calculating synchrony values, a random set of children’s data were selected as the comparator for each parent’s data so that in addition to real dyadic synchrony, “false dyadic” synchrony values (i.e., child of Dyad B paired with parent of Dyad A) would also be produced for further evaluation in later analyses. For this randomly permuted dataset, 999 samples were taken from the children’s data to match with the parent’s data. For duplicate samples, synchrony values were only calculated once for computational efficiency. NS was evaluated using the PyCWT Python package (Nedorubova et al., 2021). The continuous wavelet transform (CWT) can be used to investigate the changes in HbO at various channels and ROIs as a function of time and frequency. We used the cross-wavelet transform coherence (WTC) to investigate the concordance in these HbO changes between a parent-child dyad at task-related time points and expected frequency signatures. For each dyad, the WTC was performed using the parameters described in the PyCWT documentation, on detrended and normalized epochs of data, and on a channel-wise basis. A singularvalue was obtained for each channel by averaging the WTC over the length of the epoch, from task-related frequencies of 0.08 to 0.4 Hz (corresponding to periods of 2.5-12.5s) and excluding the cone-of-influence (COI) artifactual regions at the peripheral areas of the WTC output. We chose this frequency range because it captures low-frequency, task-related signals of interest and while providing the greatest magnitude of visibility of all areas of the WTC cone produced to visualize significant BOLD changes in fNIRS data (Nguyen et al., 2021).

The singular WTC value was calculated for each dyad contained in the randomly generated set (described above). Values were calculated across each of these dyad’s like-channels (e.g., parent’s S4_D2 to a child’s S4_D2) and in each of the 3 task conditions, unless the channel for either parent or child did not pass the HbO peak-to-peak validation, in which case that channel was excluded from analysis. We reduced this dataset by averaging the singular WTC values for the different task iterations so that, for the stress context for example, we have only one value per sampled dyad per channel, and further reduced the data by averaging channel-wise for the four identified regions-of-interest (ROIs); the left and right dorsolateral PFC (dlPFC) and the left and right ventrolateral PFC (vlPFC).

### Parent-Child Neural Synchrony Validation

Upon the completion of preprocessing, we completed permutation testing of non-concurrent pairs (real vs. false dyads) in order to verify unique concurrence due to dyadic synchrony as we have done in previous projects (Fishburn et al., 2018). This approach allowed us to confirm that the observed synchrony was driven by a dyad’s active interaction over and above the fundamental elements of shared experience between two non-partner participants (i.e., NS based on hearing the same sounds and seeing the same stimuli at the same time). NS was calculated between all possible participant pairs to determine the suitable null distribution of NS values. There were NS values for 105 real dyads and 32,448 false (null) dyads. Permutation testing was then conducted to calculate the *p*-value associated with each real dyad’s NS value by estimating the number of values from false pairings that were equal to or greater than the observed value. Constant terms were chosen to ensure that the resulting *p* values would be between zero and one. The occurrence of NS was assessed for each condition using a one-sample *t*-test. Lastly, the corresponding *p* values were corrected for multiple comparisons by calculating the Benjamini–Hochberg false discovery rate-corrected *p* value (Thissen et al., 2002) across all unique channel pairs (10 channels). Following the implementation of these t-tests, NS values were extracted from the Python-based programming environment and transferred to standard statistics software programs [Jamovi (Sahin & Aybek, 2020) and SPSS (Argyrous, 2011)].

### Analytic Strategy

To examine meaningful heterogeneity in dyadic NS in relation to key contextual and developmental variables, we applied latent profile analysis (LPA) as the person-centered analytic strategy. LPA allows for the identification of unobserved subgroups within a population based on patterns of responses across continuous variables, capturing variability that may be masked in whole-sample analyses. A series of unconditional LPA models were estimated using synchrony data to determine the most stable and interpretable number of profiles for each iteration (i.e., level) of NS analysis: (1) FC, (2) vlPFC, (3) dlPFC, and (4) left/right vlPFC and dlPFC.

A four-step approach to LPA modeling was conducted to address our study aim. First, we conducted latent profile analyses using averaged NS along the PFC – specifically in the FC, vlPFC, and dlPFC – across the three DB-DOS: BioSync task conditions, with each region examined separately. Second, we conducted a more fine-grained latent profile analysis using lateralized synchrony values from the left and right vlPFC and dlPFC, analyzed together across all three task conditions. These first two steps were used to arrive at the best-fitting unconditional LPA model prior to steps three and four. Third, we tested whether interparental conflict and perceived parental stress predict profile membership (i.e., antecedent conditional model). Finally, we examined whether children’s internalizing and externalizing symptoms differed by profile membership (i.e., distal outcome conditional model).

Model selection was guided by several fit indices: the sample-size adjusted Bayesian Information Criterion (aBIC), bootstrapped likelihood ratio test (BLRT), entropy, and minimum profile size. The aBIC is an information index used for model comparisons that considers the log-likelihood of a model, the number of parameters, and sample size, with lower values indicating better fit (Spurk et al., 2020). The BLRT is a nested model test that helps determine whether models with *k* profiles fits significantly better than *k–1* profiles; a significant p-value favors the model with more profiles, whereas a non-significant result supports the more parsimonious solution (Spurk et al., 2020). Entropy reflects how well a mixture model distinguishes between profiles, with values closer to one indicating clearer separation, though a threshold of ≥ 0.60 is generally considered acceptable (B. Muthen, 2004). Additionally, models were retained only if each profile included at least 10% of the sample (*n* = 20) to ensure stability and adequate representation (Lubke & Neale, 2006).

Next, interparental conflict and perceived parental stress were included as auxiliary variables to predict latent profile membership via multinomial logistic regression using the R3STEP approach in Mplus (Asparouhov & Muthén, 2014), which accounts for classification uncertainty and preserves the measurement model from step one. Child sex, age, race, and household socioeconomic status were included as covariates due to their established links with neurodevelopment and family stress. Differences in children’s internalizing and externalizing symptoms across latent profiles were then examined using the DU3STEP procedure, which compares distal outcome means using a Wald chi-square test. Although DU3STEP does not adjust for covariates, it preserves the latent class measurement model and accounts for classification error, allowing for valid comparisons of distal outcomes across empirically derived subgroups. LPA and subsequent auxiliary and outcome variable analyses were conducted in Mplus (v8.11; Muthen & Muthen, 2017) using full information maximum likelihood (FIML) estimation to handle missing data. SPSS (v30.0; *IBM SPSS*, n.d.) was used for data cleaning and for conducting demographic comparisons across latent profiles. This analytic sequence allowed for robust identification of latent subgroups at each level of NS analysis, accurate prediction of profile membership, and meaningful interpretation of associated developmental outcomes.

## Results

### Preliminary Analyses

Of the original 229 parent-child dyads, 194 were retained in the analytic sample after accounting for missing data on variables of interest and removing outliers in the NS data, which can bias model estimation and profile assignment in latent profile analyses. Despite some missing data, Little’s Missing Completely at Random test indicated that the data were missing completely at random, *χ²*(16) = 6.61, *p* = .98, supporting the use of FIML estimation in subsequent analyses. Two dyads were excluded as multivariate outliers based on Mahalanobis distance, using a cutoff value of *χ²*(3) = 16.27, corresponding to the 99.9^th^ percentile of the chi-squared distribution, applied to the NS data.

The analytic sample was evenly split by child sex (50% male; see Table 1). Most children were White (73%), with 27% identifying as BIPOC, and child ages ranged from 4.00 to 7.99 years old (*M* = 5.77, *SD* = 1.15). The majority of families (93%) were above the federal poverty threshold, with only 7% below. Based on the interparental conflict criteria, 54% of parents were classified as having low conflict and 46% as high conflict; among those in the high-conflict group, 73% were separated. Perceived parental stress scores ranged from 0 to 36 (*M* = 14.34, *SD* = 6.64).

**Table 1.** Demographic Characteristics of the Analytic Sample and by Latent Profile.

| Variable | Analytic Sample<br>(*n* = 194) | Latent Profile Analyses |  | Sig. |
| --- | --- | --- | --- | --- |
|  |  | Lower Baseline<br>Synchrony<br>(*n* = 132) | Higher Baseline<br>Synchrony<br>(*n* = 62) |  |
| *Child Sex* |  |  |  |  |
| Female | 97 | 73 | 24 | * |
| Male (=1) | 97 | 59 | 38 |  |
| *Child Race* |  |  |  |  |
| White | 141 | 93 | 48 | *ns* |
| BIPOC (=1) | 53 | 39 | 14 |  |
| *Child Age* | 5.77 (1.15)<br>[4.00, 7.99] | 5.73 (1.18)<br>[4.00, 7.99] | 5.86 (1.09)<br>[4.00, 7.82] | *ns* |
| *Poverty Threshold* |  |  |  |  |
| Below | 14 | 7 | 7 | *ns* |
| Above (=1) | 180 | 125 | 55 |  |
| *Interparental Conflict* |  |  |  |  |
| Low | 105 | 67 | 38 | *ns* |
| High (=1) | 89 | 65 | 24 |  |
| *Perceived Parental Stress* | 14.34 (6.64)<br>[0, 36] | 13.89 (6.45)<br>[0, 30] | 15.31 (6.97)<br>[3, 36] | *ns* |

*Note.* Values for Child Age and Perceived Parental Stress reflect Mean (Standard Deviation) [Range]. Post-hoc associations between latent profile membership and child sex, race, poverty threshold, and interparental conflict were examined using chi-square tests of independence; independent t-tests were used for child age and perceived parental stress. \* $p < .05$ , \*\* $p < .01$ , \*\*\* $p < .001$

Correlation analyses explored associations between synchrony in seven ROIs (FC, vlPFC, dlPFC, left vlPFC, left dlPFC, right vlPFC, left dlPFC) along the three task conditions and interparental conflict and perceived parental stress scores. Conflict and perceived parental stress were positively correlated (*r* = 0.31, *p* < .001), indicating that higher levels of interparental conflict were associated with greater perceived stress among parents. However, neither variable was significantly associated with neural activation in any ROI across task conditions. Among covariates, child sex, race, and age showed no significant associations with ROI activation.

Notably, families below the poverty threshold reported higher levels of interparental conflict and were more likely to have male children, as indicated by negative associations between poverty status and both conflict (*r* = –0.29, *p* < .001) and child sex (*r* = –0.17, *p* < .001). In contrast, poverty status was positively associated with right vlPFC activation during the frustration condition (*r* = 0.19, *p* < .05), suggesting greater synchrony in this region among dyads from households above the poverty line under stress.

### Global ROI LPAs: Frontal Cortex, vlPFC, dlPFC

As an initial step, we conducted a series of LPAs, separately for the FC, vlPFC, and dlPFC, using synchrony data from each of the three task conditions as indicators, to assess whether meaningful profiles emerged at each broad regional level. For the FC, the two-profile model showed a slightly lower aBIC (-3132.01) than the one-profile model (-3136.18), but low entropy (0.45) and a non-significant BLRT indicated poor class separation, favoring a one-profile solution. Similarly, for the vlPFC, although the two-profile model had a marginally lower aBIC (-2995.04) and high entropy (0.96), the BLRT was not significant, and the second profile captured only 2% of the sample (n = 4), again supporting a one-profile solution. These findings suggested that broad-region-level analyses were insufficient to capture meaningful variation in NS.

Although the dlPFC model showed somewhat better fit for the two- and three-profile solutions – reflected in decreasing aBIC values and significant BLRTs – the small size of the additional profiles (each under 10% of the sample) and only modest fit improvements limited their utility. Collectively, these results highlighted the limitations of broad regional analyses and motivated the need for a more fine-grained, disaggregated approach using lateralized ROI data. **Lateralized ROI LPAs: Left and Right vlPFC and dlPFC**

One-through three-profile solutions were estimated using 12 indicators of NS across the left and right vlPFC and dlPFC regions – capturing synchrony within each of the four regions (left vlPFC, right vlPFC, left dlPFC, right dlPFC) across the three task conditions (see Table 2). The two-profile solution yielded a lower aBIC (–10005.02) than the one-profile model (– 9984.71), and the BLRT indicated significantly improved fit from the one-to two-profile model (5017.56, *p* < .001). Entropy for the two-profile solution was 0.63, suggesting appropriate classification separation. The three-profile model provided a slightly lower aBIC (–10008.13) and improved entropy (0.69), but the BLRT was not statistically significant, suggesting the addition of a third profile did not yield a significantly better fit than the two-profile model.

**Table 2.** Predictors of Latent Profile Membership.

| **Variable** | **OR** | **SE** | **CI: L 2.5%** | **CI: U 2.5%** |
| --- | --- | --- | --- | --- |
| Interparental Conflict | 0.324* | 0.175 | 0.122 | 0.934 |
| Perceived Parental Stress | 1.071 | 0.045 | 0.986 | 1.163 |
| Child Sex (Male = 1) | 3.101* | 1.506 | 1.196 | 8.034 |
| Child Age | 1.231 | 0.245 | 0.834 | 1.817 |
| Child Race (BIPOC = 1) | 0.505 | 0.307 | 0.153 | 1.662 |
| Poverty Threshold (Above = 1) | 0.250 | 0.312 | 0.022 | 2.879 |

*Note.* The lower baseline synchrony group was used as the reference group. \* $p < .05$ , \*\* $p < .01$ ,
\*\*\* $p < .001$

Additionally, one profile in the three-profile solution comprised only 10% of the sample, raising some concerns about stability and interpretability. Given these considerations, the two-profile solution was retained as the best-fitting and most parsimonious model.

The retained two-profile solution included 132 dyads (68%) in Profile 1 and 62 dyads (32%) in Profile 2. Mean NS patterns across the two profiles differed primarily across the baseline condition, particularly in the left and right dlPFC, with one profile showing consistently higher synchrony than the other, as exhibited in Figure 1. In contrast, synchrony during the frustration and recovery conditions was comparable across profiles. Based on these patterns, the first profile is hereafter referred to as the *lower baseline synchrony* profile, and the second as the *higher baseline synchrony* profile.

**Figure 1.**
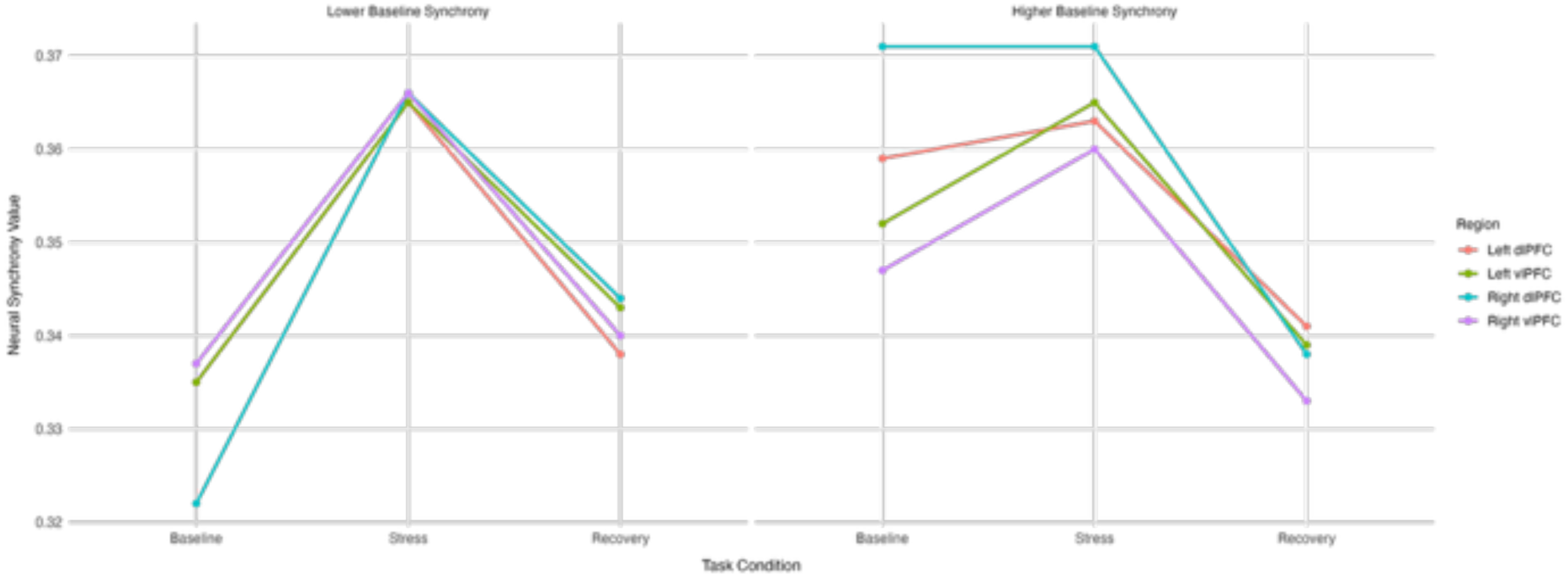
Latent profile characteristics of dyadic NS in the left and right vlPFC and dlPFC across each DB-DOS: BioSync task condition.

Descriptive analyses comparing demographic and family-level characteristics across these latent profiles using chi-square tests of independence and independent t-tests (see Table 1) indicated that child sex significantly differed by profile membership, *χ²*(1) = 4.65, *p* < 0.05.

Specifically, the higher baseline synchrony group had a greater proportion of male children (61%) relative to the lower baseline synchrony group (45%). No statistically significant differences emerged between profiles on child race, *χ²*(1) = 1.03, *p* = 0.31, poverty status, *χ²*(1) = 0.63, *p* = 0.43, or interparental conflict, *χ²*(1) = 1.89, *p* = 0.17. Similarly, there were no significant differences detected in child age, *t*(192) = –0.74, *p* = 0.46, or perceived parental stress, *t*(178) = –1.35, *p* = 0.18, between the two profiles.

### Predictors of Latent Profile Membership

Multinomial logistic regression analyses examined predictors of latent profile membership, with the lower baseline synchrony group serving as the reference. Results indicated that higher interparental conflict was significantly associated with lower odds of membership in the higher baseline synchrony profile (*OR* = 0.32, *p* < .05), suggesting that greater conflict reduces the likelihood of being in this group. Additionally, male children were over three times more likely to be classified in the higher synchrony profile compared to females (*OR* = 3.10, *p* < .05). Other predictors, including perceived parental stress, child age, race, and poverty status, were not significantly associated with profile membership (see Table 2).

### Differences in Internalizing and Externalizing Symptoms Across Latent Profiles

Significant differences emerged for internalizing symptoms across latent profiles. Children in the lower baseline synchrony group exhibited significantly higher internalizing scores (*M* = 0.27, *SE* = 0.02) compared to those in the higher baseline synchrony group (*M* = 0.18, *SE* = 0.020), *Wald χ²*(1) = 7.06, *p* < 0.01. However, no significant group differences were observed for externalizing symptoms, *Wald χ²*(1) = 1.22, *p* = 0.27, although the mean was descriptively higher in the lower synchrony group (*M* = 0.23, *SE* = 0.04) relative to the higher synchrony group (*M* = 0.12, *SE* = 0.07). These findings suggest that lower parent-child NS at baseline is selectively associated with elevated internalizing, but not externalizing, behavioral symptoms in early childhood.

## Discussion

This study applied LPA to parent-child NS data to identify distinct patterns of dyadic coordination during a mildly stressful interaction task. Building on FST and prior evidence that interparental conflict spills over into the parent-child subsystem, we examined whether synchrony profiles reflected broader family stress processes and child mental health outcomes (Cox & Paley, 1997). Although initial analyses across broad prefrontal regions showed limited differentiation, focusing on lateralized ROIs along the left and right vlPFC and dlPFC revealed two distinct profiles. These groups differed primarily in baseline synchrony during play, rather than in stress or recovery conditions, suggesting variation in foundational dyadic coordination before challenge. Importantly, higher interparental conflict predicted lower likelihood of belonging to the high baseline synchrony group, indicating that relational stress between parents may dampen neurobiological synchronization with children at the outset of interaction. Furthermore, children in the lower synchrony group reported more internalizing symptoms, highlighting NS as a potential biobehavioral pathway through which family stress shapes child emotional adjustment.

The identification of distinct synchrony profiles highlights meaningful differences in how dyads engage at a neurobiological level. Rather than viewing NS as a uniform marker of dyadic attunement and co-regulation capacities, these profiles suggest that parent-child relationships vary in the extent to which they achieve neural alignment during interaction, reflecting differences in how parents and children share attention, interpret each other’s cues, and coordinate emotional or behavioral responses (Alonso et al., 2024; Reindl et al., 2018). Prior work using the DB-DOS task has shown a consistent pattern in which NS increases between dyads from baseline to mild stress conditions, followed by a return to near-baseline levels from mild stress to recovery (e.g., Hoyniak et al., 2021; Quiñones-Camacho et al., 2022). However, in the current study, differences between profiles emerged during the baseline condition rather than during the mild stress or recovery phases, suggesting that while dyads tend to exhibit similar increases in synchrony when faced with a shared task, they differ in their initial capacity to get in sync. Such baseline differences may reflect habitual patterns of engagement shaped by broader family contexts, including interparental dynamics and everyday stress exposure (Bi et al., 2023; Leclère et al., 2014).

Given that interparental conflict, but not perceived parental stress, increased the likelihood of membership in the LB group, these patterns suggest that relational stress between parents may specifically disrupt parent-child synchrony. Notably, however, dyads in both profiles showed similar levels of synchrony during the stress and recovery conditions; differences emerged in baseline synchrony, suggesting that interparental conflict may be linked to a lagged capacity to get in sync rather than an overall inability to coordinate under challenge. Our results indicate that even when exposed to interparental conflict, some parent-child dyads maintain strong and adaptive neural alignment once engaged in a shared task, whereas others may take longer.

Accompanying these results, we unexpectedly found that child sex predicted profile membership, with male children more likely in the HB group. Although we did not have a formal hypothesis, this may reflect early differences in behavioral engagement or how parents interact with sons versus daughters. Future studies are needed to test these possibilities directly.

In addition to differences in baseline synchrony, the profiles also differed in child internalizing symptoms. Our work aligns with prior research demonstrating that reduced parent-child synchrony is associated with greater emotional difficulties in children, including anxiety and depression symptoms (Quiñones-Camacho et al., 2022; Reindl et al., 2022). Given that interparental conflict predicted membership in the LB profile, conflict may indirectly contribute to children’s internalizing problems by disrupting dyadic processes during everyday interactions at the behavioral, emotional, and neural levels. Such dyadic misalignment could limit children’s sense of felt security and their ability to effectively manage distress, potentially exacerbating internalizing symptoms over time (Figge et al., 2021; Sherrill et al., 2017). Importantly, the absence of group differences in externalizing symptoms suggests that NS may be more closely tied to inwardly directed emotional processes rather than overt behavioral dysregulation, particularly in this age group.

Though this study has notable strengths, including the novel use of LPA with NS data from a relatively large parent-child sample, several limitations warrant consideration. First, its use of cross-sectional data limits understanding of developmental change over time. Second, although our sample size was substantial for fNIRS research, some recommend at least 300 participants for optimal LPA differentiation (Asparouhov & Muthén, 2014; Nylund-Gibson et al., 2019). Third, while the sample included families from diverse backgrounds, families from impoverished or BIPOC communities were underrepresented. Moving forward, longitudinal studies with larger and more diverse samples are needed to clarify how family stress and NS shape children’s adjustment.

Overall, this study underscores the importance of examining parent-child NS as a biobehavioral mechanism linking interparental conflict to children’s emotional adjustment. By identifying distinct synchrony profiles differentiated by baseline alignment, our findings suggest that relational stress between parents may undermine foundational dyadic attunement, with implications for children’s internalizing symptoms. These results extend current research by demonstrating that interparental conflict may not uniformly disrupt parent-child coordination under stress but instead influence how readily dyads achieve synchrony during routine interactions.

